# Embrace heterogeneity to improve reproducibility: A perspective from meta-analysis of variation in preclinical research

**DOI:** 10.1101/2020.10.26.354274

**Authors:** Takuji Usui, Malcolm R. Macleod, Sarah K. McCann, Alistair M. Senior, Shinichi Nakagawa

**Author notes:** **Corresponding authors:** (TU), (AMS), (SN). These authors contributed equally to this work (Alistair M. Senior, Shinichi Nakagawa).

## Abstract

The reproducibility of research results has been a cause of increasing concern to the scientific community. The long-held belief that experimental standardization begets reproducibility has also been recently challenged, with the observation that the reduction of variability within studies can lead to idiosyncratic, lab-specific results that are irreproducible. An alternative approach is to, instead, deliberately introduce heterogeneity; known as “heterogenization” of experimental design. Here, we explore a novel perspective in the heterogenization program in a meta-analysis of variability in observed phenotypic outcomes in both control and experimental animal models of ischaemic stroke. First, by quantifying inter-individual variability across control groups we illustrate that the samount of heterogeneity in disease-state (infarct volume) differs according to methodological approach, for example, in disease-induction methods and disease models. We argue that such methods may improve reproducibility by creating diverse and representative distribution of baseline disease-state in the reference group, against which treatment efficacy is assessed. Second, we illustrate how meta-analysis can be used to simultaneously assess efficacy and stability (i.e., mean effect and among-individual variability). We identify treatments that have efficacy and are generalizable to the population level (i.e. low inter-individual variability), as well as those where there is high inter-individual variability in response; for these latter treatments translation to a clinical setting may require nuance. We argue that by embracing rather than seeking to minimise variability in phenotypic outcomes, we can motivate the shift towards heterogenization and improve both the reproducibility and generalizability of preclinical research.

## Introduction

Reproducibility of research findings – “obtaining the same results from the conduct of an independent study whose procedures are as closely matched to the original experiment as possible” [1] – is integral to scientific progress. Compelling evidence, however, suggests that irreproducibility pervades basic and preclinical research [1–5]. Moreover, animal studies motivate the development of novel treatments to be tested in clinical studies, but failure to observe effects in humans which have been reported in animal studies is commonplace [6, 7]. The conventional approach to preclinical experimental design has been to minimise heterogeneity in experimental conditions within studies to reduce the variability between animals in the observed outcomes [8]. Such rigorous standardization procedures have long been endorsed as the way to improve the reproducibility of studies by reducing within-study variability and increasing statistical power to detect treatment effects, as well as reducing the number of animals required [8, 9]. This well-established notion that standardization begets reproducibility, however, has recently been challenged.

An inadvertent consequence of standardization is that an increase in internal validity may come at the expense of external validity [10]. By reducing within-study variability, standardization may inflate between-study variability as outcomes become idiosyncratic to the particular conditions of a study, ultimately becoming only representative of local truths [10–12]. For example, in animal studies the interaction between an organism’s genotype and its local environment (i.e., phenotypic plasticity due to gene-by-environment interactions) can result in variable and discordant outcomes across laboratories using otherwise concordant methodology [13–16]. Such inconsistent outcomes may result from distinct plastic responses of animals to seemingly irrelevant and minor, unmeasured differences in environmental conditions and experimental procedures [13–18]. Through amplifying the effects of these unmeasured variables, standardization may thus weaken, rather than strengthen, reproducibility in preclinical studies.

A potential counter to this “standardization fallacy” [10] then, is to improve reproducibility by embracing, rather than minimizing, heterogeneity [10–12]. Practical solutions to enhance external validity include conducting studies across multiple laboratories to deliberately account for differences in within-lab variability [19–21], and perhaps more radically, to systematically introduce variability into experimental designs within studies [12, 22, 23]. Both simulation [11, 14, 20, 21] and empirical studies [19, 22, 24, 25] show that deliberate inclusion of more heterogeneous study samples and experimental conditions (i.e., “heterogenization”) improve external validity, and hence reproducibility, by increasing within-study (or within-lab) variability and minimizing among-study (or among-lab) variability.

Despite the promise of heterogenization, standardization remains the conventional approach in preclinical studies [26–28]. This has been partly fuelled by Russel and Birch’s [29] injunction to a “reduction in the numbers of animals used to obtain information of a given amount and precision”. Consequently, within-study variability is typically treated as a biological inconvenience that is to be minimised, rather than an outcome of interest in its own right. Embracing and quantifying heterogeneity, however, may benefit preclinical science in at least two ways. First, through comparative analyses of the variability associated with experimental procedures we may identify methodologies that introduce variation. As discussed above, by using methods that induce variation one may design a deliberately heterogeneous study with greater reproducibility [10–12]. Second, by explicitly investigating inter-individual heterogeneity in the response to drug/intervention outcomes, we may quantify the generalisability of a treatment and its translational potential. That is, a treatment with low inter-individual variation in efficacy despite heterogenization is more generalizable while a treatment with high inter-individual variation indicates the effect may be individual-specific. This may be relevant in the context of personalized medicine: A treatment associated with inter-individual variation in outcome may require tailoring in its clinical use [30]. Taking these two points together, one could argue an ideal trial would use a technical design that typically generated variation in disease state, which was then attenuated by a treatment of interest that might consistently (in all animals) or selectively (in some animals) improve outcome.

An illustrative case where the issues of reproducibility and lack of translation has been highlighted repeatedly is that of animal models of ischaemic stroke [31–33]. Several systematic reviews [34, 35] and meta-analyses [36–38] have questioned the propriety of experimental design and the choice of experimental procedures in stroke animal studies. The consequent recommendation for improving reproducibility in the field has usually been to adopt methodological procedures that minimize heterogeneity (and/or mitigate sources of bias) in phenotypic outcomes (e.g. in infarct volume or neurobehavioral outcomes) [34–38]. Furthermore, whilst potentially beneficial treatments have been identified in individual trials at the preclinical stage, intravenous thrombolysis remains the only regulatory approved treatment for ischaemic stroke [33, 39, 40]. This lack of transferable results from the preclinical to clinical stage highlights a major shortcoming for the generalizability of stroke animal models and is emblematic of translation failures generally across preclinical studies [6, 7, 33, 34].

Using the case of rat animal models of stroke as a guiding example, we highlight how recently developed methods for the meta-analysis of variation can be used to better understand biological heterogeneity. First, through analysis of variability using the log coefficient of variation (lnCV; CV representing variance relative to the mean) in control groups, we identify methodological procedures that increase variability in outcomes. Second, we show how, through the concurrent meta-analysis of mean and variance in treatment effects using the log response ratio (lnRR; i.e. ratio of means) and log coefficient of variation ratio (lnCVR), one gains additional information about the generalisability of an intervention at the individual level. Overall, we argue that the quantification of heterogeneity in phenotypic outcomes can be exploited to improve both the reproducibility and translation of animal studies.

## Results

### Dataset

We obtained data for rat animal models of ischaemic stroke from the Collaborative Approach to Meta-Analysis and Review of Animal Data from Experimental Studies (CAMARADES) database [41], focusing our meta-analysis on animal models that reported outcomes in infarct volume (see Materials and Methods for inclusion criteria of studies). We extracted data for infarct volume from 1318 control group cohorts from 778 studies for our analyses investigating the effects of methodology and variability. We extracted data for the effect of treatment on infarct volume from 1803 treatment/control group cohort pairs from 791 studies for our analyses investigating the effects of drug treatment on inter-individual variability (see S1 Data for extracted database used in this study).

### Methodology and variability

To identify methodological procedures that generated variability in disease-state, we first meta-analysed variability in infarct volume for control group animals. We quantify variability as the log coefficient of variation (lnCV) rather than the log of standard deviation because we found that our data showed a mean-variance relationship (i.e. Taylor’s Law, where the variance increases with an increase in the mean [42]; S1 Fig). Overall, the coefficient of variation (CV) in infarct volume across control groups was around 23.6% of the mean (lnCV = –1.444, CI = –1.546 to –1.342 *τ*^2^ = 0.565; Fig 1). We found large differences in variability of infarct volume (*I*^2^_*total*_ = 93.7%), suggesting that sampling variance alone cannot account for differences in the reported variability across control groups (Table 1). The *I*^2^ attributable to study was 49.6% suggesting that methodological differences across studies explained some of this heterogeneity, although a moderate amount (42.9%) of *I*^2^ remained unexplained (Table 1).

**Table 1.**
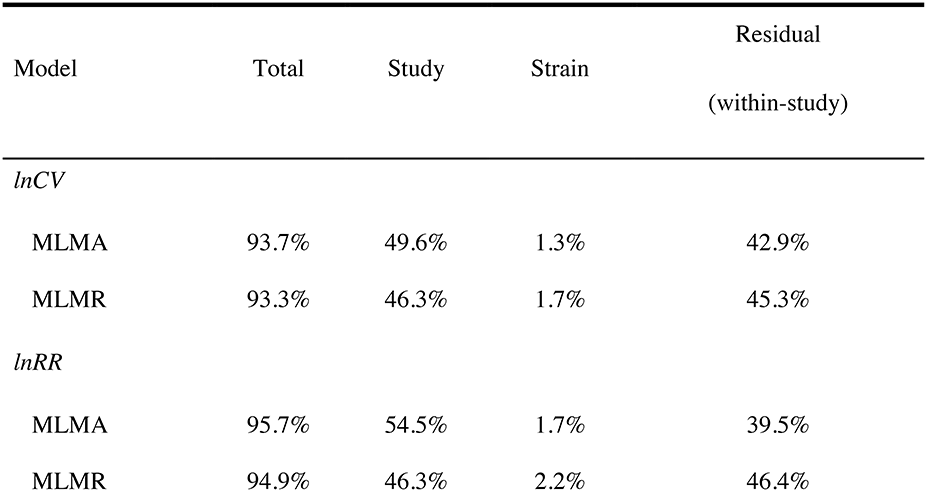
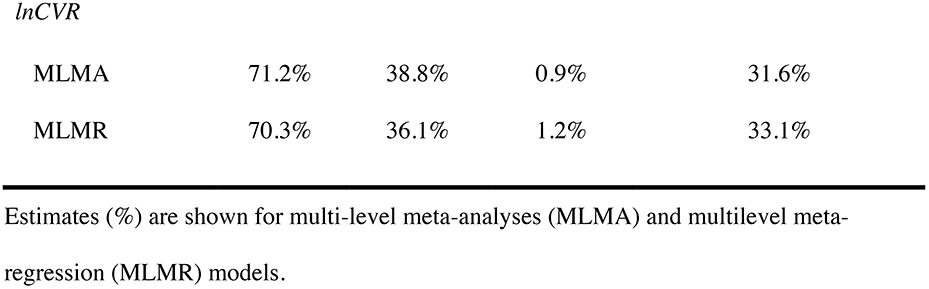
Heterogeneity (*I*^2^) estimates for analyses of methodology on variability (lnCV) and drug treatment on mean (lnRR) and variance (lnCVR) in rat infarct volume.

| Model | Total | Study | Strain | Residual<br>(within-study) |
| --- | --- | --- | --- | --- |
| *lnCV* |  |  |  |  |
| MLMA | 93.7% | 49.6% | 1.3% | 42.9% |
| MLMR | 93.3% | 46.3% | 1.7% | 45.3% |
| *lnRR* |  |  |  |  |
| MLMA | 95.7% | 54.5% | 1.7% | 39.5% |
| MLMR | 94.9% | 46.3% | 2.2% | 46.4% |
| *lnCVR* |  |  |  |  |
| MLMA | 71.2% | 38.8% | 0.9% | 31.6% |
| MLMR | 70.3% | 36.1% | 1.2% | 33.1% |

Estimates (%) are shown for multi-level meta-analyses (MLMA) and multilevel meta-regression (MLMR) models.

**Fig. 1.**
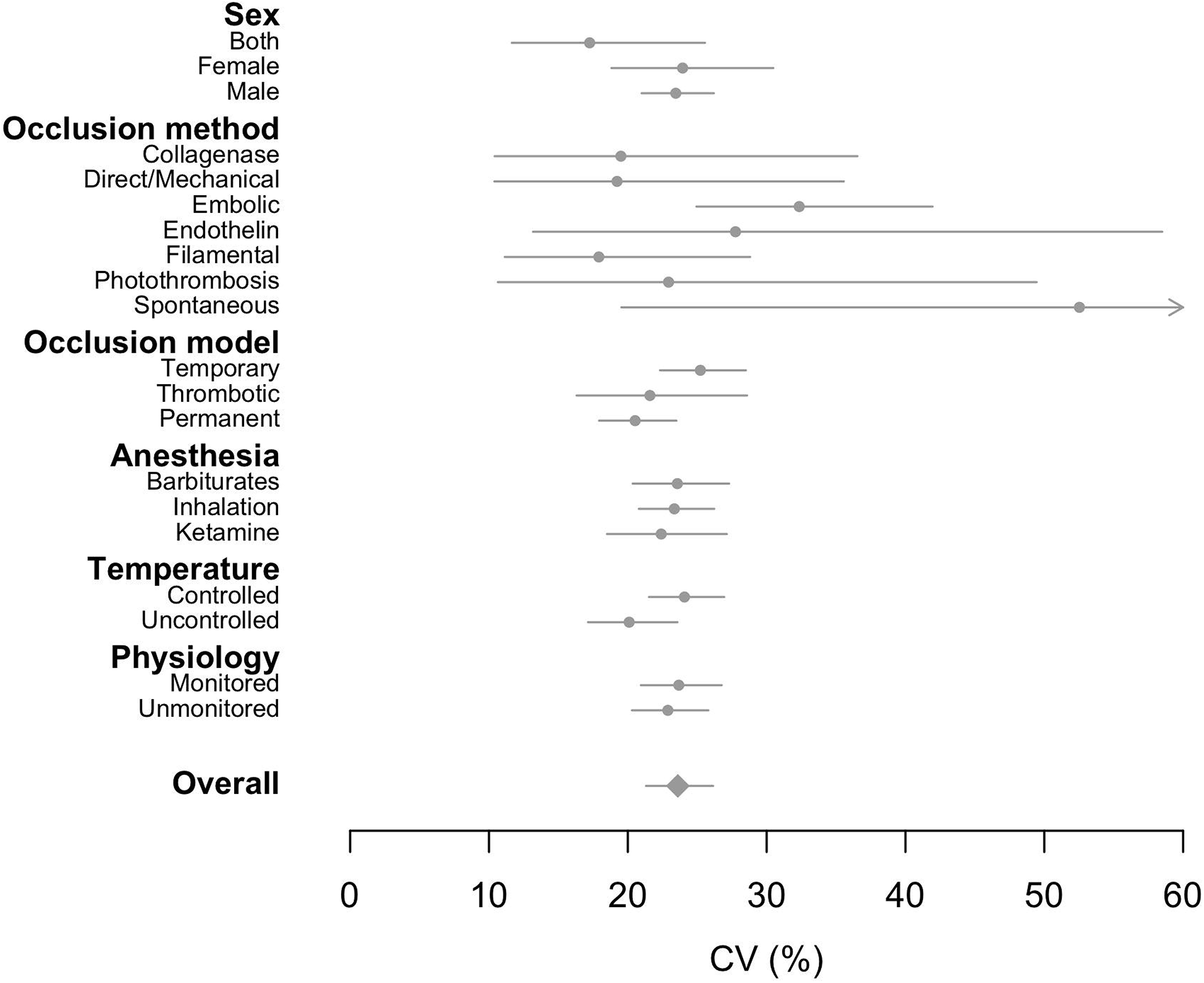
The effects of methodological parameters on variability (CV) in infarct volume across control groups. Mean estimates of unconditional (marginalized), group-specific coefficients of variation (%) are indicated as grey circles whilst the overall estimate is indicated as a grey diamond. 95% CIs are shown as grey lines and are asymmetric due to back-transformation of log coefficient of variation (lnCV) to the natural scale. Spontaneous occlusion generated the highest estimate of variability as indicated by the arrowhead. The overall and group-specific estimates were obtained from multilevel meta-analysis (MLMA) and multilevel meta-regression (MLMR) models, respectively.

We detected statistically significant differences in variability of infarct volume between various methodological approaches (Fig 1; see S1 and S2 Tables in S1 Text for unconditional and conditional model coefficients, respectively). Among occlusion methods, models with spontaneous occlusion produced the greatest variability in infarct volume (CV = 52.5%; lnCV = –0.644, –1.633 to 0.345), whilst filamental occlusion had lowest variability (CV = 17.9%; lnCV = –1.720, –2.195 to –1.244). Studies using temporary models of ischaemia had higher variability in infarct volume (CV = 25.2%; lnCV = –1.377, –1.500 to –1.255) compared to permanent models. Variability was slightly but significantly lower with longer time of damage assessment (lnCV = –1.404, –1.521 to –1.288) and greater median weight of the control group cohort (lnCV = –1.366, –1.486 to –1.245).

### Drug treatment effects and inter-individual variation

To quantify generalizability in drug treatment outcomes, we meta-analysed the mean and the coefficient of variation in infarct volume for the effects observed in control/experimental contrasts. We quantified the mean and inter-individual variability as the log response ratio (lnRR) and log coefficient of variation ratio (lnCVR), respectively. Overall, mean infarct volume in experimental groups was around 33.1%, smaller than in control groups (lnRR = –0.402, –0.461 to –0.343; Fig 2A); whilst the coefficient of variation in experimental groups was around 32.4% higher than in control groups (lnCVR = 0.280, 0.210 to 0.351; Fig 2B). Overall, heterogeneity in lnRR was very high, while that for lnCVR was moderate, and moderate amounts of heterogeneity were partitioned into the study-level for both (Table 1).

**Fig. 2.**
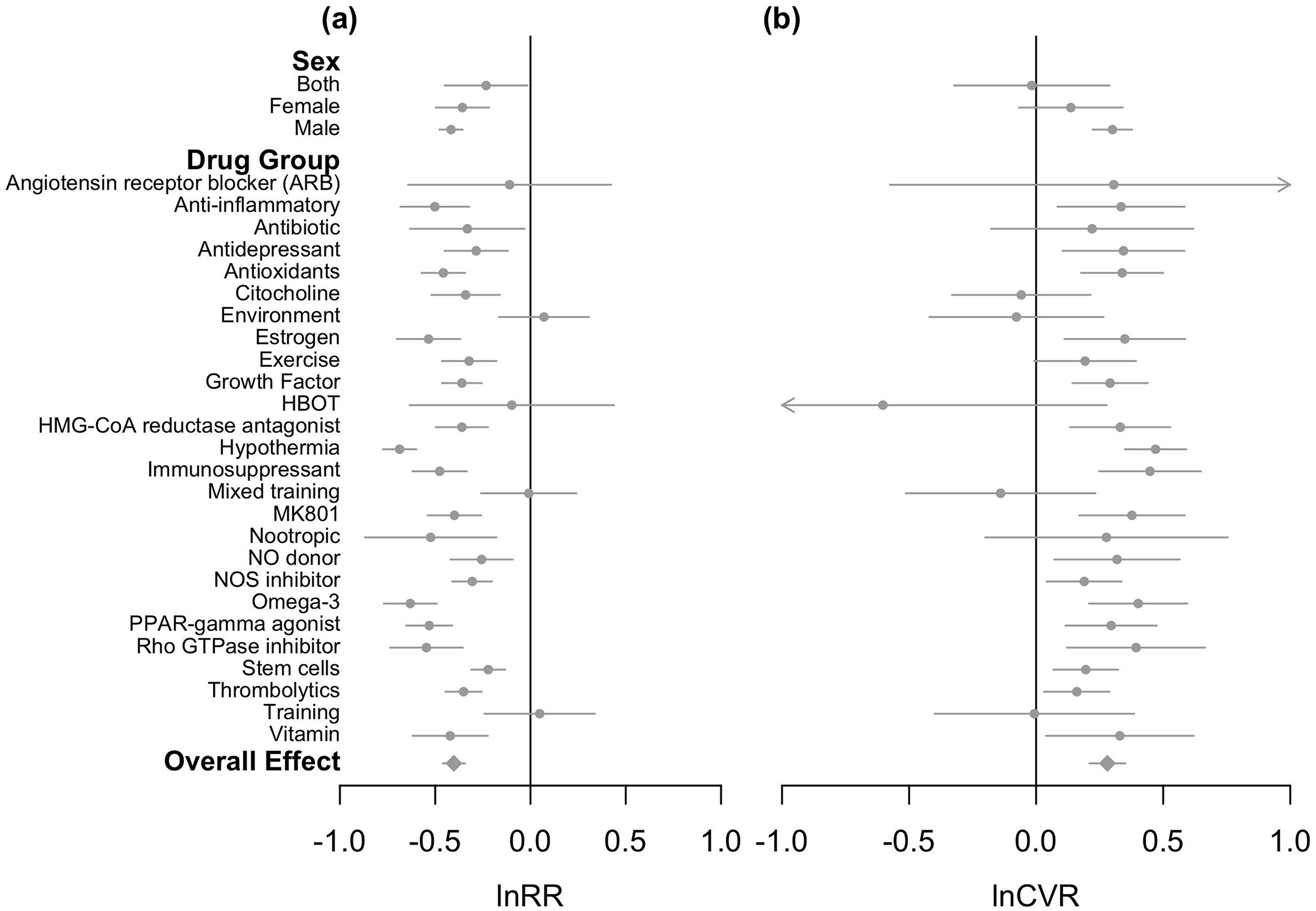
The effects of drug treatments on the difference in: (a) mean (lnRR); and (b) variability (lnCVR) in infarct volume across control and experimental rat groups. Mean estimates of unconditional (marginalized), group-specific effects are shown as grey circles whilst the overall estimate is indicated by the grey diamonds. 95% CIs are shown as grey lines. Negative lnRR estimates indicate that mean infarct volume is smaller in experimental versus control rats. Negative lnCVR estimates show that inter-individual variability in infarct volume is smaller in experimental versus control rats (e.g. HBOT indicated by left-pointing arrowhead) whilst positive lnCVR estimates show that variability in infarct volume is greater in experimental versus control rats (e.g. angiotensin receptor blockers (ARB) indicated by right-pointing arrowhead). The overall and group-specific estimates were obtained from multilevel meta-analysis (MLMA) and multilevel meta-regression (MLMR) models, respectively.

Both the mean and variability in infarct volume differed significantly across drug treatment groups (Fig 2; S3 and S4 Tables in S1 Text for unconditional and conditional model coefficients, respectively). Treatment with hypothermia resulted in the largest reduction of mean infarct volume in experimental groups relative to controls (around 49.7% lower in experimental groups than controls; lnRR = –0.687, –0.775 to –0.599). However, hypothermia also had the most variable and inconsistent effect (i.e. inter-subject variation) in reducing infarct volume, with the largest ratio of CV between experimental and control groups (inter-individual variability around 60.0% higher in experimental groups compared to controls; lnCVR = 0.470, 0.349 to 0.591). In contrast, environmental treatments were the least effective in reducing mean infarct volume (around 7.3% greater in experimental groups than controls; lnRR = 0.071, –0.166 to 0.308). Hyperbaric oxygen therapy (HBOT) has the least variable and most consistent effect on infarct volume (variability around 45.3% less in experimental groups relative to controls; lnCVR = –0.603, –1.483 to 0.277).

Thrombolytics, which include the only regulatory approved treatment (i.e., tissue plasminogen activator; tPA [42]), reduced mean infarct volume by around 29.6% in experimental relative to control groups (lnRR = –0.351, –0.446 to –0.256). The CV across experimental groups for thrombolytics was around 17.4% higher than control groups (lnCVR = 0.160, 0.031 to 0.289), but it is notable that this increased inter-subject variability is much less than that seen with hypothermia. Through quantifying variability in drug treatment outcomes, we propose that treatments be considered generalizable if they reduced mean infarct volume and concurrently show low inter-individual variability (i.e. negative lnRR and lnCVR estimates; Fig 3). Drug treatments that on average reduced infarct volume but had variable and inconsistent effects (i.e. had negative lnRR and positive lnCVR estimates; Fig 3) are ungeneralizable but might be appropriate for clinical exploitation in selected patients [30; 43]. Conversely, the least successful treatments can be identified as those that consistently do not reduce mean infarct volume (i.e. positive lnRR and lnCVR estimates; Fig 3). We explored whether the sex of groups used in experiments affected lnRR or lnCVR (see Methods for multilevel meta-regression model parameters) but differences in mean or variability of infarct volume did not vary significantly between female and male cohorts (see S5 and S6 Tables in S1 Text for contrast model estimates for sex effects).

**Fig. 3.**
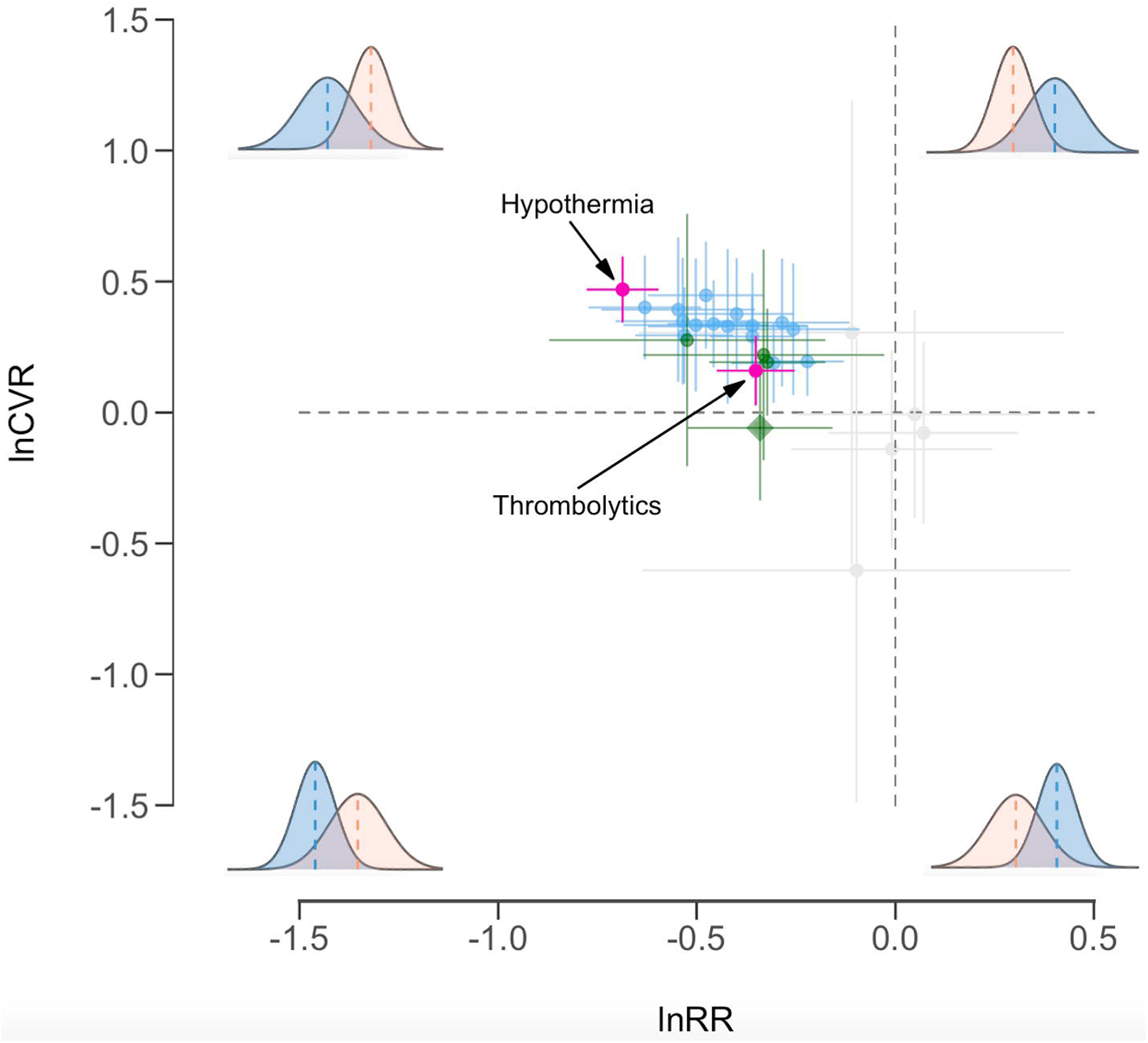
Categorization of treatment effects based on mean efficacy (lnRR) and inter-individual variability in efficacy (lnCVR). Estimates (circles) represent unconditional (marginalized), treatment-specific means (lnRR), variability (lnCVR), and their 95% CIs (solid lines) obtained from multilevel meta-regression (MLMR) models. Treatments that significantly reduce infarct volume (negative lnRR) without significantly affecting the variation are highlighted green, with citicoline indicated by a diamond as the only treatment to significantly reduce infarct volume and also have a negative point estimate of lnCVR. Treatments that significantly reduce infarct volume and increase inter-individual variability (positive lnCVR) are highlighted blue. The effects of hypothermia (most negative and positive mean and variability estimates, respectively) and thrombolytics (which include the only regulatory approved treatment) are highlighted in pink. Histograms show the relationship of the mean and variance in infarct volume between control (orange) and treatment (blue) groups in each quadrant of the graph.

## Discussion

We propose that the current failures in reproducibility and translation of preclinical trials may be due, at least in part, to the way studies are designed and assessed, which is to minimise within-study variation and ignore heterogeneity in outcomes [8, 9, 26–28]. Here, we have illustrated the potential utility of embracing such heterogeneity, through meta-analysing variability (relative variance or CV) in outcomes for rat animal models of stroke. First, by estimating the variability generated by different methodological designs applied to control animal groups, we have identified procedures that generate variability in disease-states (Fig 1). Second, we have, for the first time, quantified both the efficacy and stability (i.e., changes in the mean and variance, respectively) of stroke treatments applied to the experimental animal models (Fig 2; Fig 3), identifying potential treatments that may be generalizable versus those that require tailoring. We further discuss these results below in the context of their implications for improving the reproducibility and generalizability of preclinical studies.

### Generate variability through methodology

Among stroke animal models, studies may differ in the design of a number of parameters, including the genetic composition of animals (e.g. the sex and strain of rats used [32, 44]) as well as laboratory and operational environments (e.g. methods for stroke induction, the duration of ischemia, and the type of anaesthesia used [37, 38, 45]). However, an impediment to heterogenization is that we have not previously had reliable estimates for which methodological parameters may be most successful in generating variability in phenotypic outcomes [15]. Our results therefore quantify heterogeneity and rank the experimental factors that can generate variability in disease-state into animal models.

Our analyses of operational factors reveal that heterogeneity in outcomes may be induced by incorporating spontaneous (CV = 52.5%), embolic (CV = 32.3%), and endothelin (CV = 27.8%) methods of occlusion. Temporary models of occlusion also generate significantly more variability in disease state, than permanent models (CV = 25.2% and 20.5%, respectively). Where choices permit, we suggest that these operational design considerations are a valuable approach for introducing variability into animal models, in conjunction with more familiar proposals to diversify the laboratory environment (e.g. through differences in animal housing conditions and feeding regimens [16; 19]). Depending on the type and purpose of study, such operational and laboratory design considerations that increase heterogeneity in outcomes through environmental effects may be especially valuable when variability cannot be introduced through the animal’s genetic composition (e.g., for studies that are interested in sex-specific [46; 47] or strain-specific outcomes [44; 48]).

Our analysis is not the first to assess the effects of experimental methodology on variation in disease state in rodent models of stroke [37, 38]. Ström et al. (2013) [37] investigated similar components of experimental design on variation in infarct volume in rats. There are a number of methodological differences between their analyses and ours (e.g. differences in size of dataset and use of formal meta-analytic models). Despite these differences our quantitative results are largely concordant. Where we differ substantially is in interpretation of what is a desirable outcome. Ström et al. (2013) [37] concluded that intraluminal filament procedures are optimal as they generate minimal variation in disease outcome and maximise statistical power. Our analyses also identify that filament methods have low variation (CV = 17.9%), however, we argue that these gains in statistical power come at the cost of reduced reproducibility.

Considering genetic factors, proposals to include more heterogeneous study samples recommend the inclusion of both sexes over just male or female animals [49–51], as well as the use of multiple strains of inbred-mice and rats (or even, multiple species) [27, 52, 53]. Recent meta-analyses of variability in male and female rodents show that males may be as or more variable than females in their phenotypic response [54, 55]. We also find that male (CV = 23.5%) and female (CV = 23.9%) rats generate quantitatively equal amounts of variability, but counterintuitively find that studies that used both sexes produce the most consistent outcomes (CV = 17.3%; see S1 Table for full model coefficients). We caution that a moderate amount of the total heterogeneity remained unexplained (i.e. residual variation; Table 1), and thus these outcomes of sex on estimates of variability may be due to confounding effects of unaccounted for differences in experimental design. We therefore emphasize the importance of considering both genetic and environmental parameters for effective heterogenization of studies [56, 57].

An alternative approach to heterogenization of experimental designs within studies is to introduce variability by conducting experiments across multiple research laboratories (i.e., multi-laboratory approach) [20, 24, 58]. Importantly, such an approach inherently captures ‘unaccounted’ sources of variability in experimental conditions that are difficult to systematically manipulate within a single centre study [16, 19]. We argue that, especially where logistical constraints may hinder multi-laboratory approaches (e.g., for earlier, basic and exploratory studies), introducing heterogeneity within studies may provide the most practical alternative [23]. Indeed, by meta-analysing the variability introduced by differences in experimental methodology across studies, we can begin to find ways in which to heterogenize single studies in order to best capture the variation that exist across laboratories and studies [16; 20].

Systematically introducing variability into a system comes at the cost of reduced statistical sensitivity [8, 9] and necessitates larger studies [8, 26, 29]. These economic and ethical costs must, of course, be minimised, which can be done by identifying the most efficient means of introducing heterogeneity within experiments. It is therefore necessary to quantify the amount of variability that different experimental designs introduce, with the aim that researchers can then make informed decisions about how to most efficiently incorporate heterogeneity into study design [14–16, 20]. Identifying sources of variability through meta-analysis of variance in existing animal data as we have done here is the most practical and economic way of establishing this much needed knowledge base.

### Quantify variability to improve drug translation

Our second approach of simultaneously assessing both the mean and variation in treatment outcomes allows us to place potentially useful treatments into two, distinct categories for further exploration: 1) beneficial and generalizable interventions, which are those that consistently reduce infarct volume across individuals and; 2) beneficial but non-generalizable interventions, which on average reduce infarct volume but result in large inter-individual heterogeneity in outcomes. This latter group could even include treatments that do not necessarily reduce mean state, but have a large enough variance response to be beneficial to some [30, 43, 59].

Overall, we find that the stroke treatments in our dataset are usually effective, reducing infarct volume on average by 33.1% compared to controls. Out of these effective treatments, we identify four treatments that significantly reduced infarct volume but did not induce significant differences in the coefficient of variation across experimental and control groups (green highlights in Fig 4). Nootropic treatments reduced infarct volume on average by 40.8%, whilst citicoline, antibiotic and exercise treatments reduced infarct volume by around 27.5% to 28.8% compared to control groups. None of these treatments were estimated to significantly affect the CV, although estimated effects ranged from 5.7% smaller in experimental relative to controls for citicoline (highlighted with a triangle symbol in Fig 4), to 21.3% to 31.9% greater for the other treatments. We emphasise that these treatments may potentially be more generalizable in that the outcomes of these treatments are on average favourable, and are relatively consistent at the individual level [33, 34].

Second, we identify a handful of effective treatments that on average reduce infarct volume, but also generate significant amounts of variability in experimental groups (blue highlights in Fig 3; see S3 Table in S1 Text for rank order of unconditional estimates in mean and coefficient of variation across treatments). Of particular interest to note is that whilst thrombolytics significantly increase variability in experimental groups relative to controls, they are still relatively consistent in reducing mean infarct volume (on average reducing infarct volume by 29.6% whilst the coefficient of variation in experimental groups is only 17.4% greater than controls). Out of treatments that significantly reduce mean infarct volume, thrombolytics rank second in terms of its consistency in effect, with overlapping confidence intervals in their effects on the coefficient of variation with those of citicoline (Fig 3).

On the other hand, hypothermia is much more effective in reducing infarct volume (on average reducing infarct volume by 49.7%) but is the least consistent in doing so, estimating the greatest coefficient of variation (CV is 60.0% greater in hypothermia treated groups than concurrent controls). Interestingly, efforts to exploit hypothermia for stroke in clinical trials have so far failed to identify a patient group who might reliably benefit [60]. Other treatments that greatly reduce average infarct volume whilst increasing the variation include, for example, omega-3, rho GTPase inhibitors, and oestrogen treatments. As such, whilst these treatments confer a mean beneficial effect, this effect may not be generalizable across animals. Any future translation into clinical trials would require tailoring with effort put in to predicting response at the individual level [30]. To our knowledge, such tailoring has not been attempted because a treatment with high variability (inconsistency) is less likely to be statistically significant and pass the preclinical stage (even if it does improve a disease state) [30, 43, 59, 61]. Our study represents the first meta-analyses to quantify both the efficacy and consistency of treatment effects in animal models. We believe that this approach will forge new opportunities for improving the generalizability and translation of preclinical trials by embracing both the mean and variability in outcomes.

## Conclusion

We have demonstrated how researchers can quantitatively embrace heterogeneity in phenotypic outcomes with the aim of improving both the reproducibility and generalizability of animal models. Prior to experimentation, researchers may design their experiments by deliberately selecting methodologies that generate variability in disease-state creating a heterogenous, but broadly representative back drop of disease states against which treatment efficacy can be assessed [10–12]. Since the magnitude and direction of phenotypic expression and outcomes are determined by the interaction of genetic and environmental contexts within studies [14–16], both of these methodological factors require heterogenization in order to avoid context-specific and irreproducible outcomes across studies [16]. Post-experimentation, studies may further incorporate analyses that estimate the magnitude and direction of variability generated by treatments to identify potentially generalizable versus non-generalizable approaches. Recent meta-analyses of variability in phenotypic outcomes of animal models are beginning to illuminate the potential use of embracing different types of heterogeneities for improving reproducibility, generalizability, and translation [61–63]. We offer that comparative analyses of variability in both control and treatment groups has the potential to inform experimental design and lead to changes in both the approach and direction of follow-up studies, ultimately leading to a more successful program of reproducibility, drug discovery and translation.

## Materials and methods

### Data collection and imputation

We identified studies of rat animal models for stroke from the CAMARADES electronic database. For our analysis, we only included experimental studies that reported mean infarct volume (and their associated standard deviation and sample size) in both control and experimental groups. Where necessary we calculated the standard deviation from the standard error multiplied by the square root of (*n* – 1), where *n* is the sample size of the control or experimental group. Furthermore, when a study used multiple treatment groups for a control group, we divided the sample size of the control group equally amongst the treatment groups, which dealt with correlated errors and prevented sampling (error) variances being overly small [64]. Before calculating the effect sizes, we excluded data where: (i) the standard error was reported as zero; or (ii) the sample size of the control group when divided was equal to or less than one. We also excluded categorical predictors that were represented by fewer than five data points.

For meta-analysis of variance across methodological parameters, we focused on control groups and only included data from studies that provided sufficient group-level information on the methodology of the experiment. Specifically, we collected and coded methodological predictors as closely as possible to the predictors used by Ström et al. (2013) [37] to produce a comparable meta-analysis (see full model parameters in S1 Table in S1 Text). For meta-analysis of variance across drug treatment, we included data from studies that provided sufficient group-level information on the drug group, rat strain, and sex of experimental/control groups (see full model parameters in S3 Table in S1 Text). For all analyses, we dealt with missing data via multiple imputation [65, 66] using the package *mice* as follows: We first generated multiple, simulated datasets (*m* = 20) by replacing missing values with possible values under the assumption that data are missing at random (MAR) [66, 78]. After imputation, meta-analyses were performed on each imputed dataset (as described in *Statistical Analysis*) and model estimates were then pooled across analyses into a single set of estimates and errors.

### Calculating effect sizes

For meta-analysing variance across methodological predictors we calculated the log coefficient of variation (lnCV) and its associated sampling variance (*s*^2^_lnCV_) for each control group. Since many biological systems appear to exhibit a relationship between the mean and the variance on the natural scale (i.e., Taylor’s Law; [42, 69]), an increase in the mean may correspond to an increase in variance. Our data indeed appears to exhibit a positive relationship between log standard variation (lnSD) and log mean infarct volume (S1 Fig). When such a relationship holds in data it may be most preferable to use an effect size such as lnCV, which estimates variance accounting for the mean, and this is the approach we have taken.

For meta-analysing variance across drug treatments, we calculated the log coefficient of variance (lnCVR) and its associated sampling variance (*s*^2^_lnCVR_) as given in equations (11) and (12) in Nakagawa et al. (2015) [70] (S7 Table in S1 Text). When meta-analysing variance in the presence of Taylor’s Law as it appears in our dataset, it may be most preferable to use lnCVR (over the log variance ratio, lnVR), which gives the variance of a contrast group accounting for differences in the mean. We therefore report all results using lnCVR in the manuscript. We note, however, that both lnCV and lnCVR assumes a linear relationship between the mean and variance on the natural scale, whilst Taylor’s law states a power relationship. In addition to assessing the effects of treatments on variance, we further quantified differences in mean infarct volume by calculating the log response ratio of the mean for each control/experimental group within a study (lnRR) and its associated sampling variance (*s*^2^lnRR). For both lnRR and lnCVR we calculated effect sizes so that positive values corresponded to a larger mean or variance in the experimental group.

### Statistical analysis

We implemented multilevel meta-analytic models in a likelihood-based package using the function ‘rma.mv’ in the *metafor* package [71] as described in equation 1:

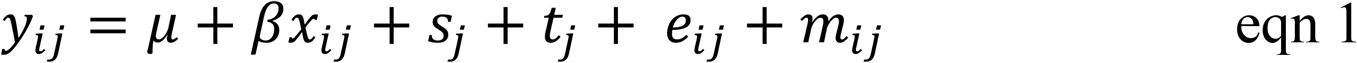

where, *y*_*ij*_ (the *i*th effect size of variability or mean infarct volume from a set of *n* effect sizes (*i* = 1, 2,…, *n*) in the *j*th study from a set of *k* studies *j* = 1, 2,…, *k*) is given by the grand mean (*μ*), the effects of fixed predictors (β*x*_*ij*_), and random effects due to study (*s*_*j*_), strain (*t*_*j*_), residual (*e*_*ij*_) and measurement error (*m*_*ij*_) for the *i*th effect size in the *j*th study. Since variability in observed effects may be explained by measurement error (*m*_*ij*_ in equation 1), we present total *I*^2^ (the percentage of variance that cannot be explained by measurement error) and study *I*^2^ (the percentage of variance explained by study-effects) to estimate the true variance in observed effects (i.e. meta-analytic heterogeneity) [72]. We interpreted *I*^2^ of 25%, 50% and 75% as small, medium, and large variance, respectively [72].

To estimate variance (lnCV) in outcome as a function of methodology in control groups we constructed two meta-analytic models. First, we fitted a multilevel meta-analysis (MLMA) with the objective of estimating the overall average variability in infarct volume across studies. MLMA included a fixed intercept and random effects described in equation 1. Second, we fitted a multilevel meta-regression (MLMR) with the objective of estimating effects of methodological predictors on variability in infarct volume, by fitting the following fixed predictors: (i) method of occlusion, (ii) sex of animal cohort, (iii) type of ischaemic model, (iv) type of anaesthetic, (v) whether experiments were temperature controlled, (vi) whether rats were physiologically monitored, (vii) mean cohort weight, and (viii) time for evaluation of damage after focal ischaemia (S1 Table in S1 Text). Mean cohort weight and time for evaluation were z-transformed prior to model fitting. We similarly constructed MLMA and MLMR models for lnRR and lnCVR (fitting each effect size as the response in separate models), to estimate the mean and variance in outcome as a function of drug treatment in our control/experimental groups, respectively. For these MLMR models, we included (i) drug treatment group, and (ii) sex of animal cohort as fixed predictors (S3 Table in S1 Text).

Fixed effects were deemed statistically significant where their 95% credible intervals (CIs) did not span zero. For interpretation of results, we back-transformed model estimates from the log to the natural scale. Finally, we tested for signs of publication bias (systematic bias in the published data due to the preferential publication of more significant results) in our data by visual inspection of funnel plots (S2 Fig) and conducting a type of Egger regression (precision-effect test and precision-effect estimate with standard errors, PET-PEESE) on lnRR [73] (see S8 Table in S1 Text for publication bias test results). Egger regression cannot be used for lnCVR, and further, it is unlikely that publication bias occurs for lnCVR because such biases are not driven by the difference in standard deviations between the experimental and control groups [74]. All meta-analyses were conducted using the ‘rma.mv’ function in the likelihood-based package *metafor* [71], on the statistical programming environment R (v 3.2.2 [75]).

## Acknowledgements

We would like to thank the CAMARADES team for help in data access and extraction, and the I-DEEL lab for providing the opportunity for TU to conduct this meta-analysis.

## Supporting information

**S1 Text.** Supporting information including tables of full model coefficients, effect size/sampling variance equations, and publication bias results. (PDF)

**S1 Fig.**
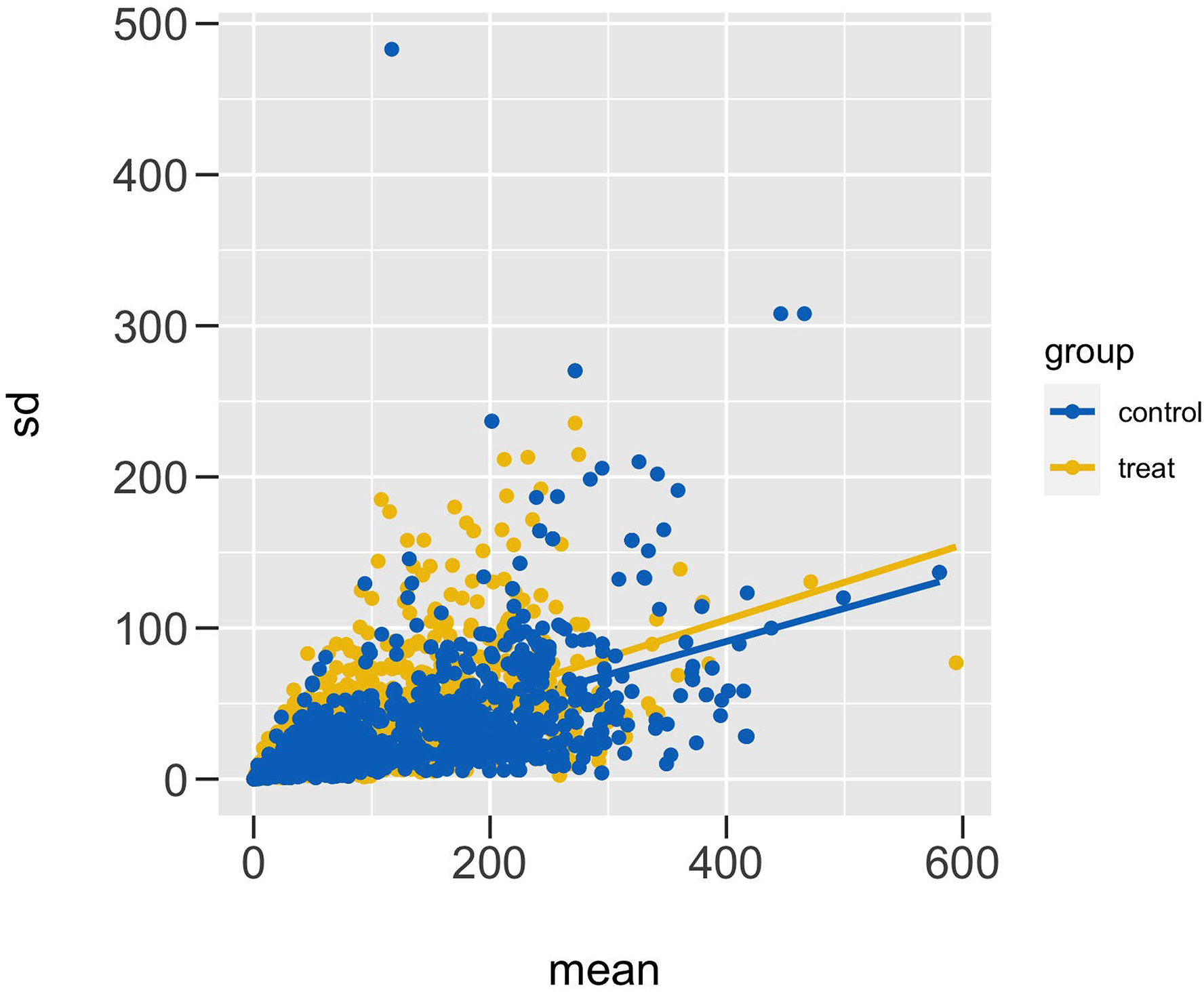
Scatter plot of mean-variance (SD) relationship in rat animal data. Point estimates for control (blue) and treatment (yellow) groups are provided, as well as their slope of linear regressions for control and experimental rat groups, respectively. Note that data points are not represented in the same units. (PDF)

**S2 Fig.**
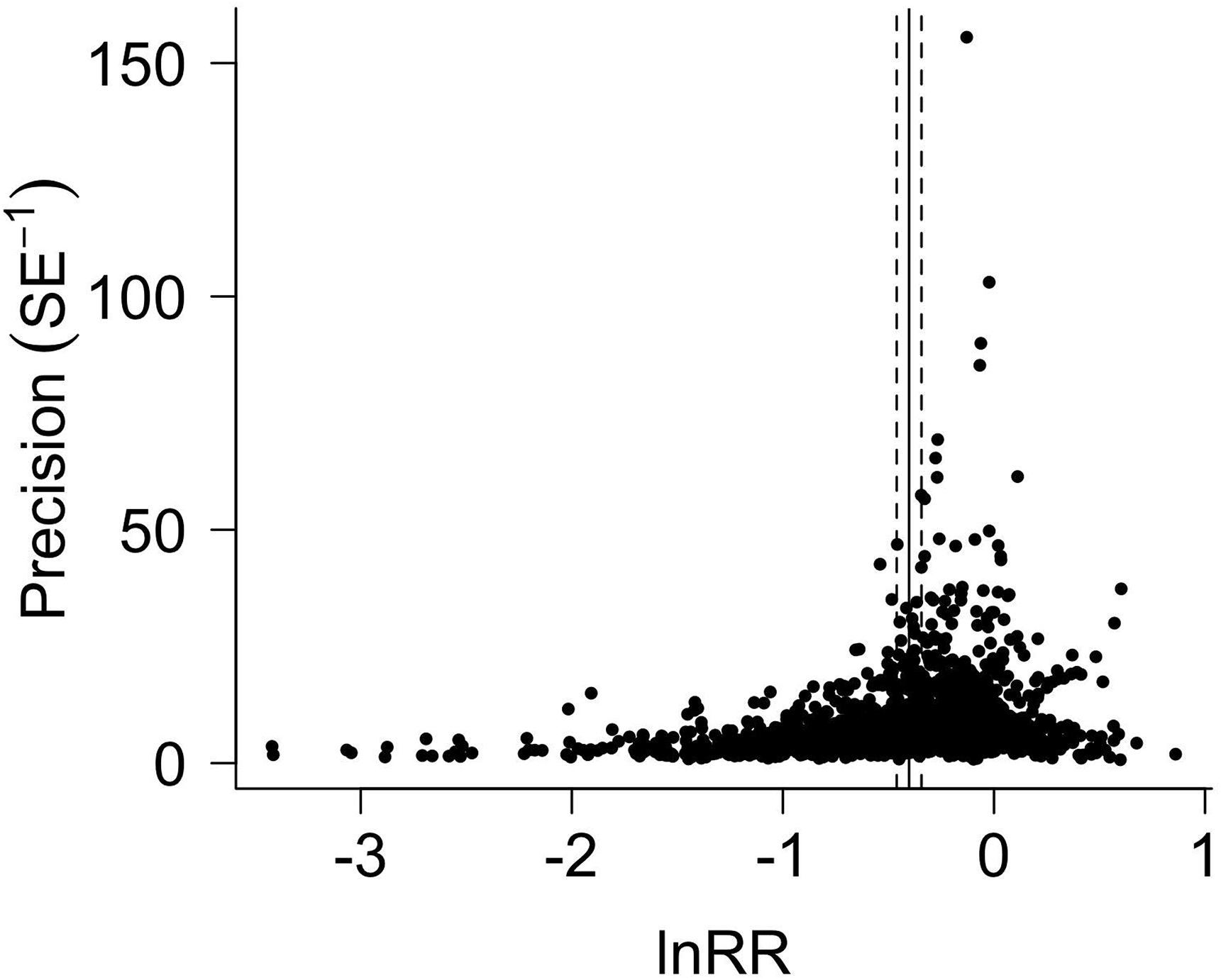
Funnel plot for log response ratio (lnRR) characterizing differences in mean infarct volume for control/treatment groups. Raw effect sizes are plotted against their precision (inverse of the square root of standard error). MLMA-model predicted mean effect size (solid vertical line) and its 95% CI (dashed lines) are shown. (PDF)

**S1 Data.** Data files for analysis of lnCV, lnRR and lnCVR in infarct volume, extracted from CAMARADES database. (RDS)

**S1 Code.** R code for conducting meta-analyses. (R-CODE)

## Author contributions

**Conceptualization:** Shinichi Nakagawa, Alistair Senior, Takuji Usui

**Data curation:** Malcolm Macleod, Sarah McCann, Takuji Usui

**Formal analysis:** Alistair Senior, Takuji Usui

**Funding acquisition:** Shinichi Nakagawa, Alistair Senior

**Supervision:** Shinichi Nakagawa, Alistair Senior

**Writing – original draft:** Takuji Usui

**Writing – review & editing:** Malcolm Macleod, Sarah McCann, Shinichi Nakagawa, Alistair Senior, Takuji Usui

