## Supplemental S1 Text for "Embrace heterogeneity to improve reproducibility: A perspective from meta-analysis of variation in preclinical research"

### S1 Supplemental Text

**S1 Table.** Unconditional (marginalized) estimates and 95% credible intervals for lnCV, obtained from multi-level regression (MLMR) models of control group infarct volume. Continuous predictors were Z-transformed prior to model fitting.

| Parameters | lnCV ($\beta$) | LCI | UCI |
|---|---|---|---|
| Sex $_{\text{BOTH}}$ | -1.757 | -2.150 | -1.364 |
| Sex $_{\text{FEMALE}}$ | -1.429 | -1.670 | -1.188 |
| Sex $_{\text{MALE}}$ | -1.450 | -1.561 | -1.339 |
| InductionMethod $_{\text{COLLAGENASE}}$ | -1.635 | -2.263 | -1.007 |
| InductionMethod $_{\text{EMBOLIC}}$ | -1.129 | -1.389 | -0.869 |
| InductionMethod $_{\text{ENDOTHELIN}}$ | -1.282 | -2.027 | -0.536 |
| InductionMethod $_{\text{FILAMENTAL}}$ | -1.720 | -2.195 | -1.244 |
| InductionMethod $_{\text{DIRECT/MECHANICAL}}$ | -1.649 | -2.264 | -1.034 |
| InductionMethod $_{\text{PHOTOTHROMBOSIS}}$ | -1.472 | -2.241 | -0.704 |
| InductionMethod $_{\text{SPONTANEOUS}}$ | -0.644 | -1.633 | 0.345 |
| IschaemiaModel $_{\text{PERMANENT}}$ | -1.583 | -1.719 | -1.448 |
| IschaemiaModel $_{\text{TEMPORARY}}$ | -1.377 | -1.500 | -1.255 |
| IschaemiaModel $_{\text{THROMBOTIC}}$ | -1.533 | -1.813 | -1.252 |
| Anesthesia $_{\text{KETAMINE}}$ | -1.496 | -1.687 | -1.304 |
| Anesthesia $_{\text{INHALATION}}$ | -1.454 | -1.570 | -1.338 |
| Anesthesia $_{\text{BARBITURATES}}$ | -1.445 | -1.592 | -1.298 |
| TemperatureControl $_{\text{NO}}$ | -1.605 | -1.764 | -1.445 |
| TemperatureControl $_{\text{YES}}$ | -1.424 | -1.536 | -1.311 |
| PhysiologyMonitored $_{\text{NO}}$ | -1.475 | -1.595 | -1.355 |
| PhysiologyMonitored $_{\text{YES}}$ | -1.441 | -1.563 | -1.318 |
| AssessTime | -1.404 | -1.521 | -1.288 |
| MidWeight | -1.366 | -1.486 | -1.245 |

**S2 Table.** Conditional estimates and 95% credible intervals for lnCV, obtained from multi-level regression (MLMR) models of control group infarct volume. Continuous predictors were Z-transformed prior to model fitting. Bold italicized estimates indicate that the 95% credible intervals do not span zero.

| Parameter | lnCV ($\beta$) | LCI | UCI |
|---|---|---|---|
| ***Intercept*** | ***-2.262*** | ***-3.024*** | ***-1.501*** |
| Sex $_{\text{FEMALE}}$ | 0.328 | -0.112 | 0.768 |
| Sex $_{\text{MALE}}$ | 0.307 | -0.075 | 0.689 |
| InductionMethod $_{\text{EMBOLIC}}$ | 0.506 | -0.169 | 1.182 |
| InductionMethod $_{\text{ENDOTHELIN}}$ | 0.500 | -0.166 | 1.167 |
| InductionMethod $_{\text{FILAMENTAL}}$ | 0.086 | -0.537 | 0.709 |
| InductionMethod $_{\text{DIRECT/MECHANICAL}}$ | 0.140 | -0.490 | 0.770 |
| InductionMethod $_{\text{PHOTOTHROMBOSIS}}$ | 0.297 | -0.393 | 0.986 |
| ***InductionMethod*** $_{\text{SPONTANEOUS}}$ | ***1.118*** | ***0.191*** | ***2.045*** |
| ***IschaemiaModel*** $_{\text{TEMPORARY}}$ | ***0.206*** | ***0.087*** | ***0.326*** |
| IschaemiaModel $_{\text{THROMBOTIC}}$ | 0.050 | -0.240 | 0.341 |
| Anesthesia $_{\text{BARBITURATES}}$ | 0.050 | -0.147 | 0.248 |
| Anesthesia $_{\text{INHALATION}}$ | 0.041 | -0.140 | 0.223 |
| ***TemperatureControl*** $_{\text{YES}}$ | ***0.181*** | ***0.041*** | ***0.321*** |
| PhysiologyMonitored $_{\text{YES}}$ | 0.034 | -0.070 | 0.138 |
| ***AssessTime*** | ***0.053*** | ***0.012*** | ***0.095*** |
| ***MidWeight*** | ***0.092*** | ***0.045*** | ***0.139*** |

**S3 Table.** Unconditional (marginalized) estimates and 95% credible intervals for lnRR and lnCVR, obtained from multi-level regression (MLMR) models of infarct volume in treatment/control groups. Treatment effects (DrugGroup) are ordered from groups that produce, on average, the greatest reduction in infarct volume (i.e. the most effective, as indicated by most negative estimates of lnRR) to groups that are, on average, the least effective.

| Parameter | lnRR | | | lnCVR | | |
|---|---|---|---|---|---|---|
| | $\beta$ | LCI | UCI | $\beta$ | LCI | UCI |
| Sex $_{\text{MALE}}$ | -0.417 | -0.478 | -0.356 | 0.300 | 0.223 | 0.378 |
| Sex $_{\text{FEMALE}}$ | -0.358 | -0.497 | -0.218 | 0.136 | -0.068 | 0.341 |
| Sex $_{\text{BOTH}}$ | -0.233 | -0.451 | -0.015 | -0.017 | -0.323 | 0.288 |
| DrugGroup $_{\text{HYPOTHERMIA}}$ | -0.687 | -0.775 | -0.599 | 0.470 | 0.349 | 0.591 |
| DrugGroup $_{\text{OMEGA-3}}$ | -0.631 | -0.770 | -0.491 | 0.402 | 0.208 | 0.595 |
| DrugGroup $_{\text{GTPase INHIBITOR}}$ | -0.546 | -0.737 | -0.355 | 0.393 | 0.122 | 0.665 |
| DrugGroup $_{\text{ESTROGEN}}$ | -0.535 | -0.702 | -0.368 | 0.349 | 0.111 | 0.587 |
| DrugGroup $_{\text{PPAR-GAMMA AGONIST}}$ | -0.531 | -0.652 | -0.410 | 0.295 | 0.115 | 0.475 |
| DrugGroup $_{\text{NOOTROPIC}}$ | -0.524 | -0.869 | -0.179 | 0.277 | -0.201 | 0.754 |
| DrugGroup $_{\text{ANTI-INFLAMMATORY}}$ | -0.502 | -0.683 | -0.322 | 0.334 | 0.084 | 0.584 |
| DrugGroup $_{\text{IMMUNOSUPPRESSANT}}$ | -0.477 | -0.619 | -0.334 | 0.448 | 0.248 | 0.649 |
| DrugGroup $_{\text{ANTIOXIDANT}}$ | -0.457 | -0.571 | -0.343 | 0.339 | 0.177 | 0.501 |
| DrugGroup $_{\text{VITAMIN}}$ | -0.422 | -0.620 | -0.224 | 0.329 | 0.038 | 0.620 |
| DrugGroup $_{\text{MK801}}$ | -0.399 | -0.540 | -0.258 | 0.377 | 0.169 | 0.585 |
| DrugGroup $_{\text{GROWTH FACTOR}}$ | -0.360 | -0.464 | -0.256 | 0.291 | 0.143 | 0.439 |
| DrugGroup $_{\text{HMG-CoA REDUCTASE ANTAGONIST}}$ | -0.360 | -0.497 | -0.222 | 0.331 | 0.134 | 0.529 |
| DrugGroup $_{\text{THROMBOLYTICS}}$ | -0.351 | -0.446 | -0.256 | 0.160 | 0.031 | 0.289 |
| DrugGroup $_{\text{CITOCHOLINE}}$ | -0.340 | -0.520 | -0.161 | -0.058 | -0.332 | 0.215 |
| DrugGroup $_{\text{ANTIBIOTIC}}$ | -0.331 | -0.632 | -0.031 | 0.220 | -0.178 | 0.619 |
| DrugGroup $_{\text{EXERCISE}}$ | -0.322 | -0.466 | -0.179 | 0.193 | -0.008 | 0.393 |

| | | | | | | |
|---|---|---|---|---|---|---|
| DrugGroup NOS INHIBITOR | -0.306 | -0.411 | -0.202 | 0.189 | 0.041 | 0.337 |
| DrugGroup ANTIDEPRESSANT | -0.285 | -0.451 | -0.118 | 0.344 | 0.104 | 0.584 |
| DrugGroup NO DONOR | -0.257 | -0.420 | -0.093 | 0.318 | 0.071 | 0.565 |
| DrugGroup STEM CELLS | -0.222 | -0.311 | -0.132 | 0.196 | 0.068 | 0.323 |
| DrugGroup ANGIOTENSIN RECEPTOR BLOCKER (ARB) | -0.110 | -0.642 | 0.423 | 0.306 | -0.575 | 1.187 |
| DrugGroup HBOT | -0.098 | -0.634 | 0.438 | -0.603 | -1.483 | 0.277 |
| DrugGroup MIXED TRAINING | -0.009 | -0.259 | 0.241 | -0.140 | -0.513 | 0.234 |
| DrugGroup TRAINING | 0.048 | -0.242 | 0.338 | -0.007 | -0.399 | 0.386 |
| DrugGroup ENVIRONMENT | 0.071 | -0.166 | 0.308 | -0.077 | -0.420 | 0.265 |

**S4 Table.** Conditional estimates and 95% credible intervals for lnRR and lnCVR, obtained from multi-level regression (MLMR) models of infarct volume in treatment/control groups. Bold italicized estimates indicate that the 95% credible intervals do not span zero.

| **Parameter** | | **lnRR** | | | **lnCVR** | |
|---|---|---|---|---|---|---|
| | *β* | LCI | UCI | *β* | LCI | UCI |
| Intercept | ***-0.695*** | ***-0.783*** | ***-0.607*** | ***0.488*** | ***0.367*** | ***0.609*** |
| Sex BOTH | 0.184 | -0.031 | 0.399 | ***-0.318*** | ***-0.623*** | ***-0.013*** |
| Sex FEMALE | 0.059 | -0.079 | 0.197 | -0.164 | -0.369 | 0.042 |
| DrugGroup ANGIOTENSIN RECEPTOR BLOCKER (ARB) | ***0.577*** | ***0.043*** | ***1.111*** | -0.164 | -1.048 | 0.720 |
| DrugGroup ANTI-INFLAMMATORY | 0.185 | -0.002 | 0.371 | -0.136 | -0.398 | 0.127 |
| DrugGroup ANTIBIOTIC | ***0.355*** | ***0.052*** | ***0.658*** | -0.250 | -0.654 | 0.155 |
| DrugGroup ANTIDEPRESSANT | ***0.402*** | ***0.230*** | ***0.574*** | -0.126 | -0.378 | 0.126 |
| DrugGroup ANTIOXIDANT | ***0.229*** | ***0.107*** | ***0.351*** | -0.131 | -0.311 | 0.048 |
| DrugGroup CITOCHOLINE | ***0.346*** | ***0.160*** | ***0.533*** | ***-0.528*** | ***-0.813*** | ***-0.244*** |
| DrugGroup ENVIRONMENT | ***0.758*** | ***0.517*** | ***0.998*** | ***-0.547*** | ***-0.899*** | ***-0.196*** |
| DrugGroup ESTROGEN | 0.152 | -0.023 | 0.326 | -0.121 | -0.375 | 0.133 |
| DrugGroup EXERCISE | ***0.365*** | ***0.215*** | ***0.514*** | ***-0.277*** | ***-0.491*** | ***-0.063*** |
| DrugGroup GROWTH FACTOR | ***0.327*** | ***0.214*** | ***0.439*** | ***-0.179*** | ***-0.345*** | ***-0.013*** |
| DrugGroup HBOT | ***0.589*** | ***0.050*** | ***1.128*** | ***-1.073*** | ***-1.958*** | ***-0.188*** |
| DrugGroup HMG-CoA REDUCTASE ANTAGONIST | ***0.327*** | ***0.183*** | ***0.471*** | -0.139 | -0.350 | 0.073 |
| DrugGroup IMMUNOSUPPRESSANT | ***0.210*** | ***0.062*** | ***0.358*** | -0.022 | -0.236 | 0.192 |
| DrugGroup MIXED TRAINING | ***0.678*** | ***0.420*** | ***0.935*** | ***-0.610*** | ***-0.995*** | ***-0.224*** |
| DrugGroup MK801 | ***0.288*** | ***0.140*** | ***0.435*** | -0.093 | -0.315 | 0.129 |
| DrugGroup NOOTROPIC | 0.163 | -0.185 | 0.511 | -0.193 | -0.677 | 0.291 |
| DrugGroup NO DONOR | ***0.430*** | ***0.260*** | ***0.600*** | -0.152 | -0.411 | 0.108 |
| DrugGroup NOS INHIBITOR | ***0.381*** | ***0.265*** | ***0.496*** | ***-0.281*** | ***-0.450*** | ***-0.111*** |
| DrugGroup OMEGA-3 | 0.056 | -0.088 | 0.201 | -0.068 | -0.275 | 0.138 |

| | | | | | | |
|---|---|---|---|---|---|---|
| DrugGroup PPAR-GAMMA AGONIST | ***0.156*** | ***0.026*** | ***0.285*** | -0.175 | -0.371 | 0.022 |
| DrugGroup GTPase INHIBITOR | 0.140 | -0.055 | 0.336 | -0.077 | -0.358 | 0.205 |
| DrugGroup STEM CELLS | ***0.465*** | ***0.366*** | ***0.564*** | ***-0.274*** | ***-0.424*** | ***-0.125*** |
| DrugGroup THROMBOLYTICS | ***0.336*** | ***0.233*** | ***0.438*** | ***-0.310*** | ***-0.458*** | ***-0.162*** |
| DrugGroup TRAINING | ***0.735*** | ***0.442*** | ***1.028*** | ***-0.477*** | ***-0.877*** | ***-0.077*** |
| DrugGroup VITAMIN | ***0.265*** | ***0.061*** | ***0.468*** | -0.141 | -0.443 | 0.162 |

**S5 Table.** Conditional estimates and 95% credible intervals for lnRR and lnCVR, obtained from contrast multi-level regression (MLMR) models to assess the effect of sex on infarct volume. The intercept here represents studies in which "Both" sexes were used. Bold italicized estimates indicate that the 95% credible intervals do not span zero.

| Parameters | lnRR | | | lnCVR | | |
|---|---|---|---|---|---|---|
| | $\beta$ | LCI | UCI | $\beta$ | LCI | UCI |
| Intercept | ***-0.511*** | ***-0.740*** | ***-0.282*** | 0.171 | -0.153 | 0.494 |
| Sex FEMALE | -0.125 | -0.352 | 0.103 | 0.154 | -0.174 | 0.482 |
| Sex MALE | -0.184 | -0.399 | 0.031 | ***0.318*** | ***0.013*** | ***0.623*** |
| DrugGroup ANGIOTENSIN RECEPTOR BLOCKER (ARB) | ***0.577*** | ***0.043*** | ***1.111*** | -0.164 | -1.048 | 0.720 |
| DrugGroup ANTI-INFLAMMATORY | 0.185 | -0.002 | 0.371 | -0.136 | -0.398 | 0.127 |
| DrugGroup ANTIBIOTIC | ***0.355*** | ***0.052*** | ***0.658*** | -0.250 | -0.654 | 0.155 |
| DrugGroup ANTIDEPRESSANT | ***0.402*** | ***0.230*** | ***0.574*** | -0.126 | -0.378 | 0.126 |
| DrugGroup ANTIOXIDANT | ***0.229*** | ***0.107*** | ***0.351*** | -0.131 | -0.311 | 0.048 |
| DrugGroup CITOCHOLINE | ***0.346*** | ***0.160*** | ***0.533*** | ***-0.528*** | ***-0.813*** | ***-0.244*** |
| DrugGroup ENVIRONMENT | ***0.758*** | ***0.517*** | ***0.998*** | ***-0.547*** | ***-0.899*** | ***-0.196*** |
| DrugGroup ESTROGEN | 0.152 | -0.023 | 0.326 | -0.121 | -0.375 | 0.133 |
| DrugGroup EXERCISE | ***0.365*** | ***0.215*** | ***0.514*** | ***-0.277*** | ***-0.491*** | ***-0.063*** |
| DrugGroup GROWTH FACTOR | ***0.327*** | ***0.214*** | ***0.439*** | ***-0.179*** | ***-0.345*** | ***-0.013*** |
| DrugGroup HBOT | ***0.589*** | ***0.050*** | ***1.128*** | ***-1.073*** | ***-1.958*** | ***-0.188*** |
| DrugGroup HMG-CoA REDUCTASE ANTAGONIST | ***0.327*** | ***0.183*** | ***0.471*** | -0.139 | -0.350 | 0.073 |
| DrugGroup IMMUNOSUPPRESSANT | ***0.210*** | ***0.062*** | ***0.358*** | -0.022 | -0.236 | 0.192 |
| DrugGroup MIXED TRAINING | ***0.678*** | ***0.420*** | ***0.935*** | ***-0.610*** | ***-0.995*** | ***-0.224*** |
| DrugGroup MK801 | ***0.288*** | ***0.140*** | ***0.435*** | -0.093 | -0.315 | 0.129 |
| DrugGroup NOOTROPIC | 0.163 | -0.185 | 0.511 | -0.193 | -0.677 | 0.291 |
| DrugGroup NO DONOR | ***0.430*** | ***0.260*** | ***0.600*** | -0.152 | -0.411 | 0.108 |
| DrugGroup NOS INHIBITOR | ***0.381*** | ***0.265*** | ***0.496*** | ***-0.281*** | ***-0.450*** | ***-0.111*** |

| | | | | | | |
|---|---|---|---|---|---|---|
| DrugGroup $_{\text{OMEGA-3}}$ | 0.056 | -0.088 | 0.201 | -0.068 | -0.275 | 0.138 |
| DrugGroup $_{\text{PPAR-GAMMA AGONIST}}$ | ***0.156*** | ***0.026*** | ***0.285*** | -0.175 | -0.371 | 0.022 |
| DrugGroup $_{\text{GTPase INHIBITOR}}$ | 0.140 | -0.055 | 0.336 | -0.077 | -0.358 | 0.205 |
| DrugGroup $_{\text{STEM CELLS}}$ | ***0.465*** | ***0.366*** | ***0.564*** | ***-0.274*** | ***-0.424*** | ***-0.125*** |
| DrugGroup $_{\text{THROMBOLYTICS}}$ | ***0.336*** | ***0.233*** | ***0.438*** | ***-0.310*** | ***-0.458*** | ***-0.162*** |
| DrugGroup $_{\text{TRAINING}}$ | ***0.735*** | ***0.442*** | ***1.028*** | ***-0.477*** | ***-0.877*** | ***-0.077*** |
| DrugGroup $_{\text{VITAMIN}}$ | ***0.265*** | ***0.061*** | ***0.468*** | -0.141 | -0.443 | 0.162 |

**S6 Table.** Conditional estimates and 95% credible intervals for lnRR and lnCVR, obtained from contrast multi-level regression (MLMR) models to assess the effect of sex on infarct volume. The intercept here represents studies in which only "Female" sex was used. Bold italicized estimates indicate that the 95% credible intervals do not span zero.

| Parameters | lnRR | | | lnCVR | | |
|---|---|---|---|---|---|---|
| | $\beta$ | LCI | UCI | $\beta$ | LCI | UCI |
| Intercept | **-0.636** | **-0.795** | **-0.476** | **0.324** | **0.091** | **0.558** |
| Sex $_{\text{BOTH}}$ | 0.125 | -0.103 | 0.352 | -0.154 | -0.482 | 0.174 |
| Sex $_{\text{MALE}}$ | -0.059 | -0.197 | 0.079 | 0.164 | -0.042 | 0.369 |
| DrugGroup $_{\text{ANGIOTENSIN RECEPTOR BLOCKER (ARB)}}$ | ***0.577*** | ***0.043*** | ***1.111*** | -0.164 | -1.048 | 0.720 |
| DrugGroup $_{\text{ANTI-INFLAMMATORY}}$ | 0.185 | -0.002 | 0.371 | -0.136 | -0.398 | 0.127 |
| DrugGroup $_{\text{ANTIBIOTIC}}$ | ***0.355*** | ***0.052*** | ***0.658*** | -0.250 | -0.654 | 0.155 |
| DrugGroup $_{\text{ANTIDEPRESSANT}}$ | ***0.402*** | ***0.230*** | ***0.574*** | -0.126 | -0.378 | 0.126 |
| DrugGroup $_{\text{ANTIOXIDANT}}$ | ***0.229*** | ***0.107*** | ***0.351*** | -0.131 | -0.311 | 0.048 |
| DrugGroup $_{\text{CITOCHOLINE}}$ | ***0.346*** | ***0.160*** | ***0.533*** | ***-0.528*** | ***-0.813*** | ***-0.244*** |
| DrugGroup $_{\text{ENVIRONMENT}}$ | ***0.758*** | ***0.517*** | ***0.998*** | ***-0.547*** | ***-0.899*** | ***-0.196*** |
| DrugGroup $_{\text{ESTROGEN}}$ | 0.152 | -0.023 | 0.326 | -0.121 | -0.375 | 0.133 |
| DrugGroup $_{\text{EXERCISE}}$ | ***0.365*** | ***0.215*** | ***0.514*** | ***-0.277*** | ***-0.491*** | ***-0.063*** |
| DrugGroup $_{\text{GROWTH FACTOR}}$ | ***0.327*** | ***0.214*** | ***0.439*** | ***-0.179*** | ***-0.345*** | ***-0.013*** |
| DrugGroup $_{\text{HBOT}}$ | ***0.589*** | ***0.050*** | ***1.128*** | ***-1.073*** | ***-1.958*** | ***-0.188*** |
| DrugGroup $_{\text{HMG-CoA REDUCTASE ANTAGONIST}}$ | ***0.327*** | ***0.183*** | ***0.471*** | -0.139 | -0.350 | 0.073 |
| DrugGroup $_{\text{IMMUNOSUPPRESSANT}}$ | ***0.210*** | ***0.062*** | ***0.358*** | -0.022 | -0.236 | 0.192 |
| DrugGroup $_{\text{MIXED TRAINING}}$ | ***0.678*** | ***0.420*** | ***0.935*** | ***-0.610*** | ***-0.995*** | ***-0.224*** |
| DrugGroup $_{\text{MK801}}$ | ***0.288*** | ***0.140*** | ***0.435*** | -0.093 | -0.315 | 0.129 |
| DrugGroup $_{\text{NOOTROPIC}}$ | 0.163 | -0.185 | 0.511 | -0.193 | -0.677 | 0.291 |
| DrugGroup $_{\text{NO DONOR}}$ | ***0.430*** | ***0.260*** | ***0.600*** | -0.152 | -0.411 | 0.108 |
| DrugGroup $_{\text{NOS INHIBITOR}}$ | ***0.381*** | ***0.265*** | ***0.496*** | ***-0.281*** | ***-0.450*** | ***-0.111*** |

| | | | | | | |
|---|---|---|---|---|---|---|
| DrugGroup OMEGA-3 | 0.056 | -0.088 | 0.201 | -0.068 | -0.275 | 0.138 |
| DrugGroup PPAR-GAMMA AGONIST | ***0.156*** | ***0.026*** | ***0.285*** | -0.175 | -0.371 | 0.022 |
| DrugGroup GTPase INHIBITOR | 0.140 | -0.055 | 0.336 | -0.077 | -0.358 | 0.205 |
| DrugGroup STEM CELLS | ***0.465*** | ***0.366*** | ***0.564*** | ***-0.274*** | ***-0.424*** | ***-0.125*** |
| DrugGroup THROMBOLYTICS | ***0.336*** | ***0.233*** | ***0.438*** | ***-0.310*** | ***-0.458*** | ***-0.162*** |
| DrugGroup TRAINING | ***0.735*** | ***0.442*** | ***1.028*** | ***-0.477*** | ***-0.877*** | ***-0.077*** |
| DrugGroup VITAMIN | ***0.265*** | ***0.061*** | ***0.468*** | -0.141 | -0.443 | 0.162 |

**S7 Table.** Effect sizes and sampling variances used in meta-analysis of variance (a) across methodological predictors and (b) across drug treatment groups. Equations and the model type in which the effect size was used are also given. $\bar{x}$ and s is the mean and SD of the group infarct volume, $n$ is the sample size, CV is the coefficient of variation, and $\rho$ is the correlation between the mean and standard deviation on the log scale. Subscripts C and E refer to control and treatment groups, respectively.

| Effect Size | Outcome Measure | Equation | Sampling Variance ($s^2_{\text{Effect Size}}$) | Model |
|---|---|---|---|---|
| | | **(a) Meta-analysis of methodological predictors** | | |
| lnCV | Coefficient of Variation | $\ln\left(\frac{s}{\bar{x}}\right) + \frac{1}{2(n-1)}$ | $\frac{s^2}{n\bar{x}^2} + \frac{1}{2(n-1)} - 2\rho\sqrt{\frac{s^2}{n\bar{x}^2}\frac{1}{2(n_C-1)}}$ | Arm-based / Meta-regression |
| | | **(b) Meta-analysis of drug treatment** | | |
| lnRR | Mean | $\ln\left(\frac{\bar{x}_E}{\bar{x}_C}\right)$ | $\frac{s_C^2}{n_C\bar{x}_C^2} + \frac{s_E^2}{n_E\bar{x}_E^2}$ | Contrast-based |
| lnCVR | Coefficient of Variance | $\ln\left(\frac{CV_E}{CV_C}\right) + \frac{1}{2(n_E-1)} - \frac{1}{2(n_C-1)}$ | $\frac{s_C^2}{n_C\bar{x}_C^2} + \frac{1}{2(n_C-1)} - 2\rho\sqrt{\frac{s_C^2}{n_C\bar{x}_C^2}\frac{1}{2(n_C-1)}} + \frac{s_E^2}{n_E\bar{x}_E^2} + \frac{1}{2(n_E-1)} - 2\rho\sqrt{\frac{s_E^2}{n_E\bar{x}_E^2}\frac{1}{2(n_E-1)}}$ | Contrast-based |

**S8 Table.** Results from Egger regression (PET-PEESE) on lnRR to test for publication bias. This procedure fits the square-root of sampling variance as a moderator (slope estimate and 95% credible intervals shown in the first half of the table). If this estimate is significant, we then fit the sampling variance (second half of the table). The intercept from this latter model indicates a 'potentially' bias-corrected, modified meta-analytic mean. In our case, the biased-corrected estimate is a 28.0% decline, compared with the original estimate without correction, which is 29.6% decline. These values indicate that although this analysis detected a sign of publication bias, the effect of this bias is very small (1.6% difference). Bold italicized estimates indicate that the 95% credible intervals do not span zero.

| **Parameter** | **lnRR ($\beta$)** | **LCI** | **UCI** |
|---|---|---|---|
| ***Intercept*** | ***-0.1525*** | ***-0.2102*** | ***-0.0949*** |
| **sqrt(sampling variance)** | ***-1.5185*** | ***-1.7131*** | ***-1.3239*** |
| ***Intercept*** | ***-0.3285*** | ***-0.3813*** | ***-0.2757*** |
| ***sampling variance*** | ***-1.8964*** | ***-2.2141*** | ***-1.5787*** |
